# Predicting rifampicin resistance in *M. tuberculosis* using machine learning informed by protein structural and chemical features

**DOI:** 10.1101/2024.08.15.608097

**Authors:** Charlotte I Lynch, Dylan Adlard, Philip W Fowler

**Affiliations:** Nuffield Department of Medicine, University of Oxford, Oxford, UK; National Institute of Health Research Oxford Biomedical Research Centre, John Radcliffe Hospital, Headley Way, Oxford, UK; Health Protection Research Unit in Healthcare Associated Infections and Antimicrobial Resistance, University of Oxford, UK

**Keywords:** Tuberculosis, machine learning, rifampicin, genetics, antimicrobial resistance, diagnostics

## Abstract

**Background:** Rifampicin remains a key antibiotic in the treatment of tuberculosis. Despite advances in cataloguing resistance-associated variants (RAVs), novel and rare mutations in the relevent gene, *rpoB*, will be encountered in clinical samples, complicating the task of using genetics to predict whether a sample is resistant or not to rifampicin. We have trained a series of machine learning models with the aim of complementing genetics-based drug susceptibility testing.

**Methods:** We built a Test+Train dataset comprising 219 susceptible mutations and 46 RAVs. Features derived from the structure of the RNA polymerase or the change in chemistry introduced by the mutation were considered, however, only a few, notably the distance from the rifampicin binding site, were found to be predictive on their own. Due to the paucity of RAVs we used Monte Carlo cross-validation with 50 repeats to train four different machine learning models.

**Results:** All four models behaved similarly with sensitivities and specificities in the range 0.84-0.88 and 0.94-0.97 although we preferred the ensemble of Decision Tree models as they are easy to inspect and understand. We showed that measuring distances from molecular dynamics simulations did not improve performance.

**Conclusions:** It is possible to predict whether a mutation in *rpoB* confers resistance to rifampicin using a machine learning model trained on a combination of structural, chemical and evolutionary features, however performance is moderate and training is complicated by the lack of data.

## Introduction

Antimicrobial resistance is a growing global problem. This is a particular issue in tuberculosis, a disease which is often fatal if untreated. In 2022, 10.6 million people developed tuberculosis (TB) and 1.3 million died [1]. Tuberculosis is difficult to treat and requires multiple antibiotics to be taken for months. Drug susceptibility testing is therefore key for both effective treatment and the prevention of the spread of drug resistance. Whilst culture-based testing remains the “gold standard” it is slow, expensive and requires considerable expertise. Nucleic acid amplification tests (NAAT), such as the Cepheid GeneXpert MTB/RIF^®^, take a matter of hours and have been heavily subsidised, encouraging their uptake. GeneXpert MTB/RIF detects nucleotide mutations in a small region of the gene which is the target for rifampicin [2–4]. Any mutation is assumed to confer resistance to rifampicin, and since isoniazid resistance is epidemiologically associated with rifampicin resistance, resistance to isoniazid is then inferred. For many years multi-drug resistance (MDR) tuberculosis has been defined as resistance to both these drugs. Since NAAT tests predominate, most of the c. 410,000 cases of drug-resistant TB reported in 2022 were resistant to rifampicin [1] and are likely to be resistant to isoniazid and possibly other antibiotics as well.

Whole genome sequencing (WGS) and related methods, such as targeted next generation sequencing (tNGS), have the potential to offer much greater resolution since they can detect resistance in a panel of drugs. This has been spurred on in recent years by the release of catalogues of mutations in *M. tuberculosis*, the aetiological agent of TB, by the WHO [5–7]. Like NAATs, these approaches are inferential and so suffer from the weakness that they cannot return a result when a novel or rare mutation is detected in a gene known to be associated with resistance.

Machine-learning (ML) models can potentially address this weakness, thereby complementing WGS-based approaches. Previous ML studies have typically focussed on genetic features, with some also including lineage, geographic, and structural features [8–10], utilizing a range of model architectures. Zhang *et al* [11]. and Kuang *et al*. [12] trained convolutional neural networks (CNNs) to predict resistance to various TB drugs, while Chowdhury *et al*. [13] have employed more traditional ML models incorporating structural information to predict mycobacterial capreomycin resistance. The advantage of using structural features is that they potentially contain information about the physical mechanism of drug inhibition, rather than abstracting it away as genetic features do. Carter *et al*. [14] and Karmakar *et al*. [15] both predicted pyrazinamide resistance using structurally-informed models, and notably, Portelli *et al*. [16] have trained a range of ML models on structural features to predict rifampicin resistance due to mutations within *rpoB*.

Rifampicin inhibits bacterial replication by sterically occluding the extension of 2-nt to 3-nt mRNA in the RpoB subunit of the RNA polymerase (Fig. 1a) [17]. Consequently, resistance-associated variants (RAVs) within *rpoB* are predominantly located in the so-called “rifampicin resistance defining region” (RRDR), which forms the rifampicin binding interface. This contrasts with variants associated with pyrazinamide resistance, for example, which are more even spatially distributed in its target protein, PncA [14]. We hypothesize that this well-defined mechanism of resistance should facilitate efficient label discrimination, enabling accurate predictions with fewer features and a less complex model.

**Figure 1:**
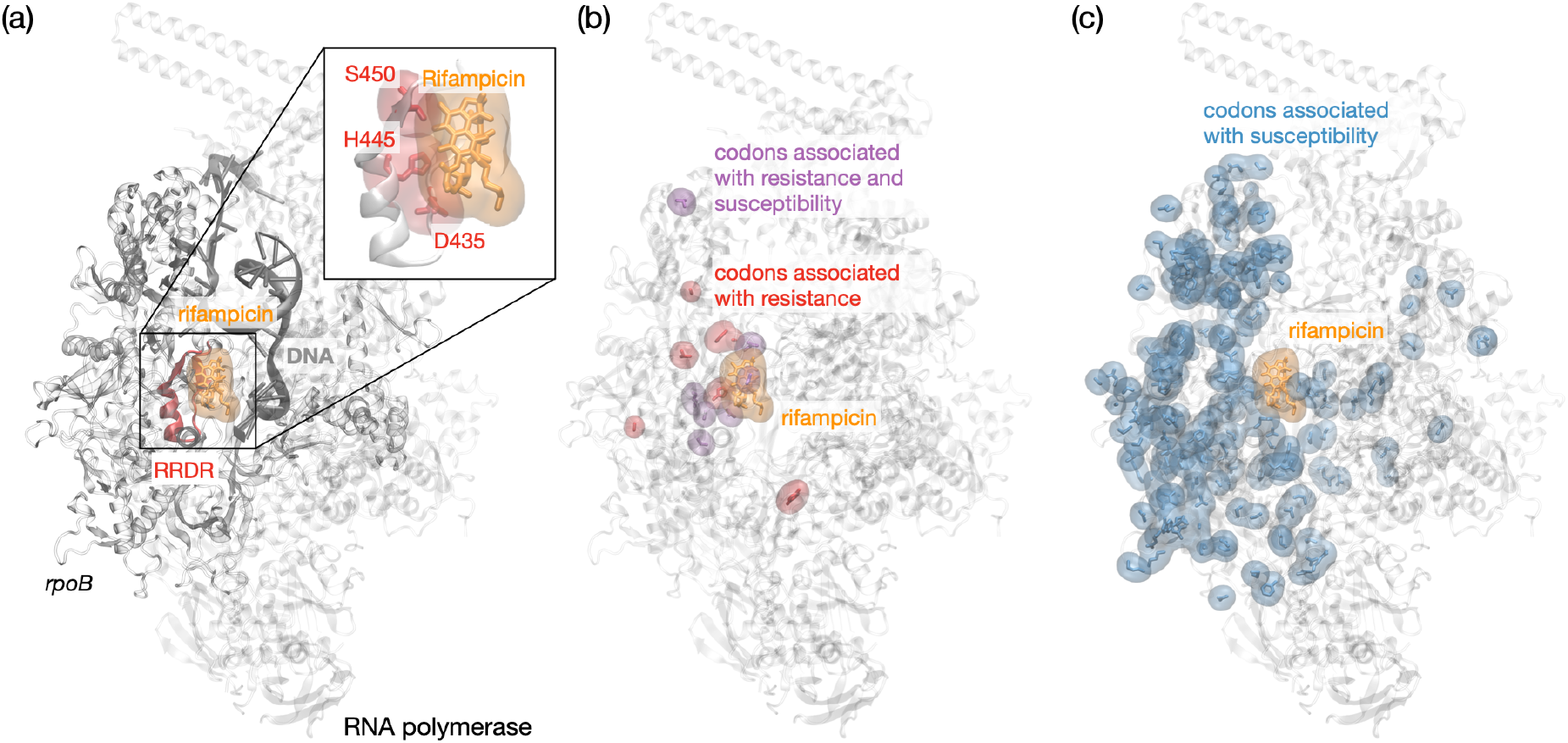
Rifampicin binds to the RNA polymerase (RNAP) and prevents the extension of nascent RNA. (a) The majority of resistance-associated variants (RAVs) occur within the so-called rifampicin-resistance determining region (RRDR, red) located in the RpoB subunit of the RNA polymerase (PDB 5UH6, [17]). The three most common residues associated with resistance are Ser450, His445 and Asp435. (b) The majority of RAVs are located close to the rifampicin binding site. (c) Mutations not associated with resistance occur throughout the RpoB protein subunit.

In this paper, we shall determine how well a series of machine learning models can predict whether missense mutations in the *rpoB* gene result in susceptibility or instead confer resistance to rifampicin. Our models are all trained on the same Train+Test dataset, and are informed by a combination of protein structural (e.g. distances), chemical and evolutionary features to attempt to capture as much information as possible on how the drug binds to the protein. Since proteins are dynamic under physiological conditions, we also investigate if using distances derived from classical molecular simulations, rather than from the experimental structure, improves performance.

## Materials and Methods

### Dataset

We built a dataset comprising unique missense mutations derived from 33,091 isolates collected by the CRyPTIC consortium [18]. Each isolate underwent whole genome sequencing (WGS) using Illumina sequencing and rifampicin susceptibility testing. The consensus genome for each sample was inferred from the short genetic read files using the minos variant caller as incorporated into v0.12.4 of the clockwork pipeline [19, 20]. The resulting variant call (VCF) files were then processed by gnomonicus (v2.5.1) [21] which calculated all relevant genetic variants, including translating the sequence to identify amino acid mutations. Filtering to only include samples with missense mutations in the *rpoB* gene resulted in 15,356 samples.

Binary resistant/susceptibility results were produced using a wide range of antibiotic susceptibility testing (AST) methods. The two most common were Mycobacterial Growth Indicator Tube (MGIT) and 96-well broth microdilution plate. The former is standard, whilst the latter was developed for the CRyPTIC project [20]. Due to the challenges associated with interpreting Mycobacterial growth in broth microdilution plates, we only included minimum inhibitory concentrations (MIC) where at least two independent reading methods concurred, thereby minimizing measurement error [22, 23]. A research epidemiological cutoff was then applied to classify each measurement as either resistant or susceptible [20]. In total this resulted in 14,523 samples, of which 10,058 were resistant and 4,465 were susceptible.

### Train+Test dataset

To generate a Train+Test dataset of mutations with labels, we constructed a catalogue using the binary phenotypes of the samples. The underlying algorithm was inspired by methodologies employed by Walker et al. [24] and the first and second editions of the WHO catalogue of RAVs [5, 6], where the definitive defectives algorithm [25] was used to identify and classify benign variants. Under the assumption that resistance-associated variants (RAVs) in *rpoB* will always induce a resistant phenotype, and to capture all possible RAVs, we used a one-tailed binomial test against an arbitrary 5% background rate. This test was conducted under the null hypothesis at 95% confidence that there is no statistically significant difference between the proportion of resistance in samples containing the mutation, when it is the only mutation present in *rpoB*, and the background rate. If the null hypothesis was accepted, the mutation was classified as susceptible (S); if rejected, it was classified as resistant (R). This approach generated a catalogue of 357 mutations. However, after selecting missense mutations that occur within the non-attenuated region of the experimental structure, the Train+Test dataset comprised 219 susceptible mutations and only 46 resistant mutations.

Best practice would be, at this point, to set aside a fraction of the dataset to use for validation, however this is not practical here due to the paucity of mutations associated with resistance (positives). Additionally, the presence of even one or two false negatives would have a spuriously large impact on the apparent performance. Instead we opted to perform Monte Carlo cross-validation [26] using an 80:20 split whereby for each iteration 80% of the dataset is used to train the models and the performance is calculated using the remaining 20%. We arbitrarily repeat this 50 times, each time using a different (known and specified) random seed to ensure reproducibility.

### Training and hyperparameter tuning

We chose to train four different machine learning models using the Python3 scikit-learn library [27]: these were logistic regression (LR), decision tree (DT), random forest (RF), and gradient-boosted decision tree (GBDT) algorithms. Features were scaled, decision thresholds selected, and model hyperparameters tuned via grid searches under 5-fold stratified cross-fold validation on the Train dataset within each iteration. Given the clinical aim of minimising false negatives (Very Major Errors – VMEs), all models were optimized for recall. Sensitivity (i.e. recall) and specificity measured on the test set in each iteration were used as the primary performance metrics throughout the study: these equate to one minus the VME or Major Error rate, respectively.

### Determination of structural and chemical features

A range of features were generated for each mutation using the Python package sbmlcore [28] from an experimental structure of the *M. tuberculosis* RNA polymerase with rifampicin bound, resolved to 3.8 Å [17]. Structural features included the distances from the C_*α*_ atom of the residue in question to the centres of mass of the rifampicin and mRNA molecules, antisense and sense DNA strands and magnesium and zinc ions, *φ* and *ψ* protein backbone angles (all calculated using MDAnalysis [29, 30]), protein secondary structure (using STRIDE) [31], the change in the number of hydrogen bond donors and acceptors, and the change in the solvent accessible surface area (SASA, using FreeSASA) [32]. Chemical features included changes on mutation in molecular weight, volume, hydropathy, isoelectric point, and a chemical similarity score [33]. Additionally, we included changes in a score designed to predict whether mutations are neutral or deleterious (SNAP2) [34], as well as the results of two machine learning models that predict the change in protein stability due to a mutation (DeepDDG [35] and RasP [36]). Finally, to assess the effect of dynamics on the measured distances, we measured the minimum distances from a series of published molecular dynamics simulations of the RNAP protein [37]. To avoid outliers, the minimum distance was defined as the 5th percentile of the distances aggregated from all three simulations.

## Results

### Spatial distribution of mutations

Despite *rpoB* being an essential gene, we observe missense mutations along the entire length of the gene. Of the 46 unique resistance-associated variants, 36 are located within the RRDR (codons 426– 452), with the remainder (Fig. 1b) close to the rifampicin binding site. In contrast, only 13 of the 219 susceptible mutations are found within the RRDR and the remainder (Fig. 1c) appear scattered throughout the structure of RpoB. We accordingly hypothesised that distance-based features would be predictive.

### A few features are very predictive

The correlation between the chosen features (Fig. S1) shows that many features, as expected, are confounded. This is particularly true for distances, but also applies to features representing stability (SNAP2 score) and residue flexibility (temperature factors) due to the relatively buried nature of the RRDR. We examined the predictive power of each feature by training a univariate logistic regression model using Monte Carlo cross-validation with 50 repeats. The performance of these models (Fig. 2) clearly indicate that only a handful of features possess any significant predictive power in isolation – notably, any of the distances, SNAP2 score and the structural temperature factor.

**Figure 2:**
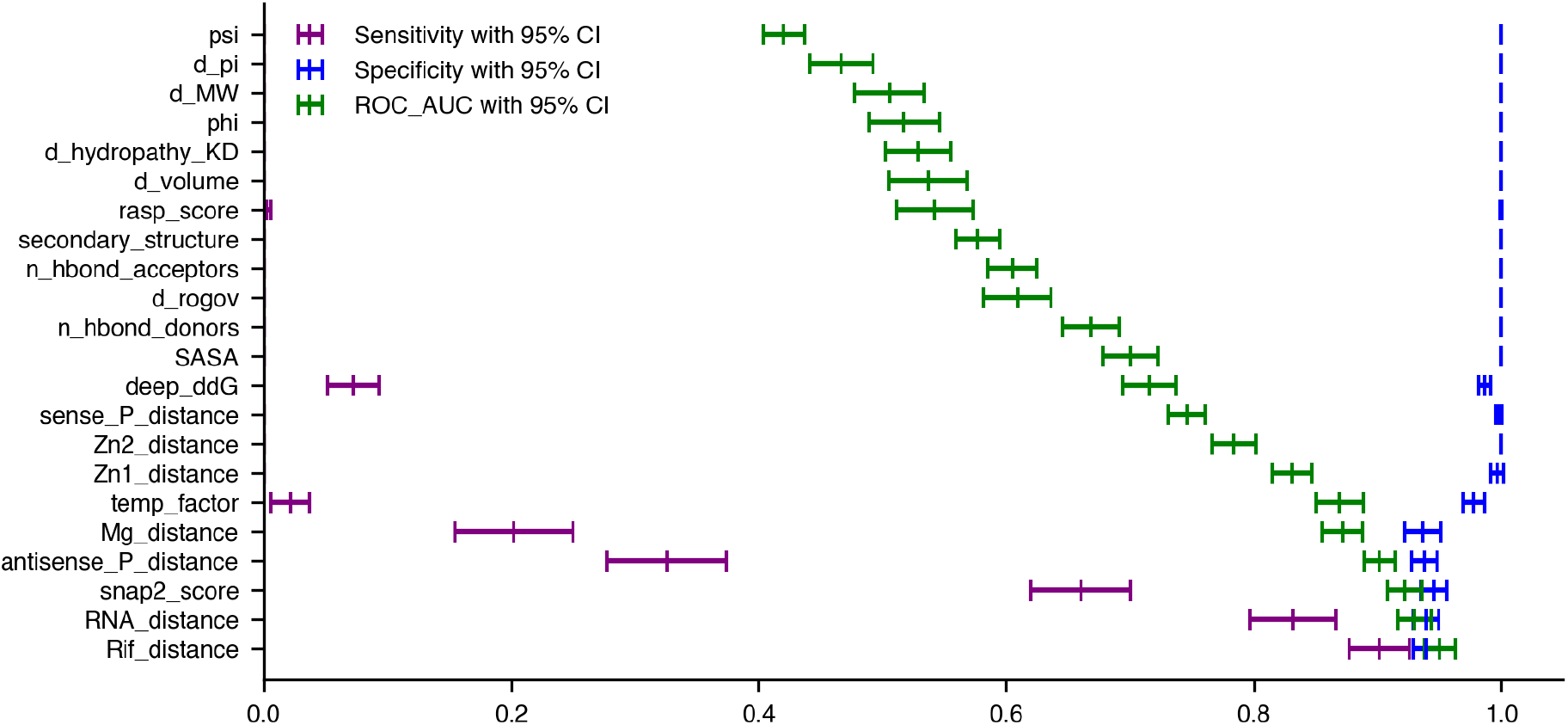
(a) Sensitivity, specificity, and area under the ROC curve were calculated from Monte Carlo cross-validated univariate logistic regression models, each trained on a single feature in isolation, using 50-repeat Monte Carlo cross-validation, and ordered by ROC AUC.

To determine how many predictive features are required for a stable model, we performed backwards elimination of features as ordered according to the receiver operator characteristic area under the curve (ROC AUC) values of the univariate Logistic Regression models (Fig. 2). This demonstrated one could only train on distance to rifampicin and still get robust performance (Fig. 3). Since several of these features are likely to interact, we chose to retain the top four most discriminatory features: these are the distances to the centres of mass of rifampicin and mRNA, the SNAP2 score, and the temperature factors, whilst noting that these features are confounded to some degree.

**Figure 3:**
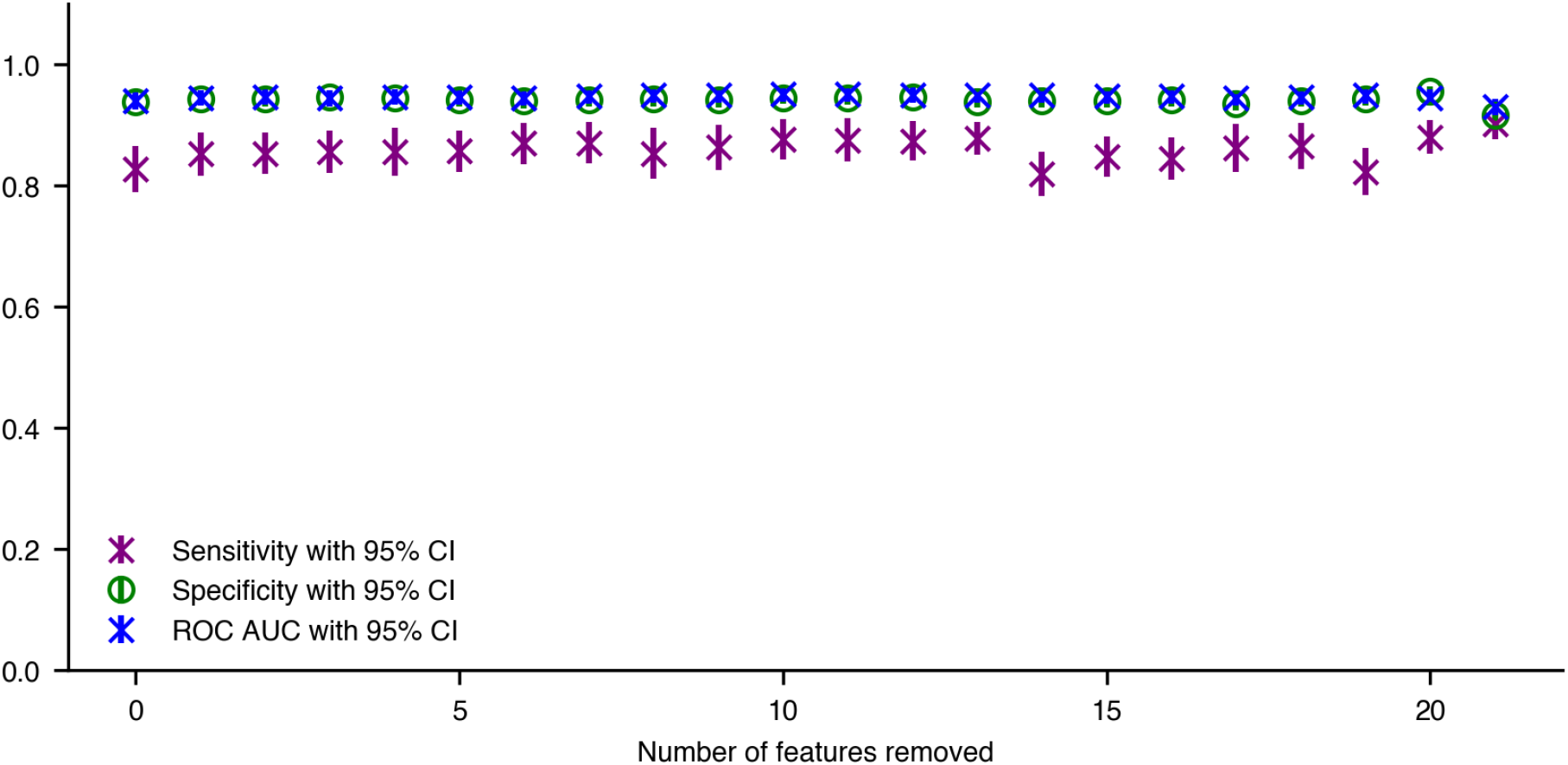
Mean sensitivity, specificity, and area under the ROC curve calculated for each backwards elimination step, with SEM error bars generated under 50-repeat Monte Carlo cross-validation.

### All models predict rifampicin resistance with moderate sensitivity

Following feature selection and hyperparameter tuning, four machine learning models (logistic regression, decision tree, random forest, and gradient boosted decision tree) were trained using the above four features and validated using Monte Carlo cross-validation with 50 iterations (Fig. 4, Table 1). All models performed similarly well in the aggregate, with mean sensitivity and specificity values in the ranges of 0.84-0.88 and 0.94-0.97, respectively. The variation in performance within each iteration was also consistent across models. Given the clear relationship between resistance and proximity to rifampicin, the simplicity and transparency of the decision tree (sensitivity 0.876 *±* 0.028, specificity 0.96 *±* 0.008) is particularly attractive, as it can provide effective, interpretable decisions (Fig. S2a). All subsequent analysis is therefore done with the ensemble of decision tree models.

**Table 1:** Mean sensitivity, specificity, and area under the receiver operating characteristic curve (ROC) with standard error of the mean 95% confidence intervals, for the logistic regression (LR), decision tree (DT), random forest (RF), and gradient boosted decision tree (GBDT) models, as well as for the decision tree trained on structural data calculated from molecular dynamics trajectories (Dynamic DT), all trained and tested on the same data under a 50-repeat Monte Carlo cross validation protocol.

|  | Sensitivity | Specificity | Area under ROC |
| --- | --- | --- | --- |
| **LR** | $0.836 \pm 0.040$ | $0.941 \pm 0.011$ | $0.949 \pm 0.013$ |
| **DT** | $0.876 \pm 0.028$ | $0.963 \pm 0.008$ | $0.915 \pm 0.018$ |
| **RF** | $0.842 \pm 0.032$ | $0.965 \pm 0.008$ | $0.942 \pm 0.012$ |
| **GBDT** | $0.856 \pm 0.030$ | $0.961 \pm 0.009$ | $0.938 \pm 0.011$ |
| **Dynamic DT** | $0.835 \pm 0.033$ | $0.956 \pm 0.009$ | $0.895 \pm 0.018$ |

**Figure 4:**
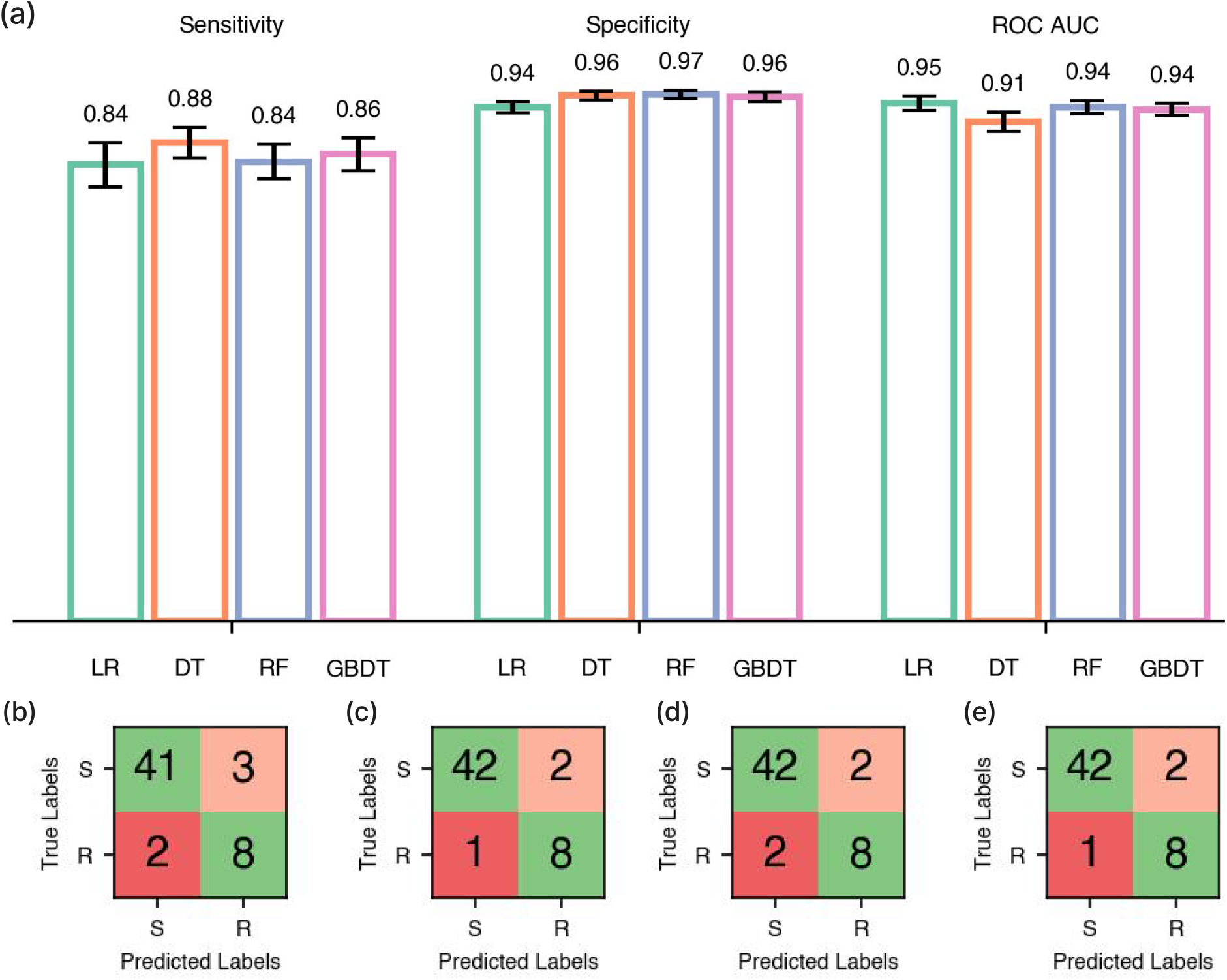
(a) Mean sensitivity, specificity, and area under the ROC curve values for hyperparameter-tuned Logistic Regression (LR), Decision Tree (DT), Random Forest (RF), and Gradient Boosted Decision Tree (GBDT) models trained under a 50-repeat Monte Carlo cross-validation protocol with 80:20 Train:Test splits. Standard error of the mean 95% confidence intervals are overlaid. All models were trained on distance to rifampicin, distance to mRNA, SNAP2 score, and temperature factors. (b-e) Confusion matrices showing the mean predicted True Positive, False Positive, True Negative, and False Negative values for the Test sets generated in (a) for each model in the same order.

**Figure 5:**
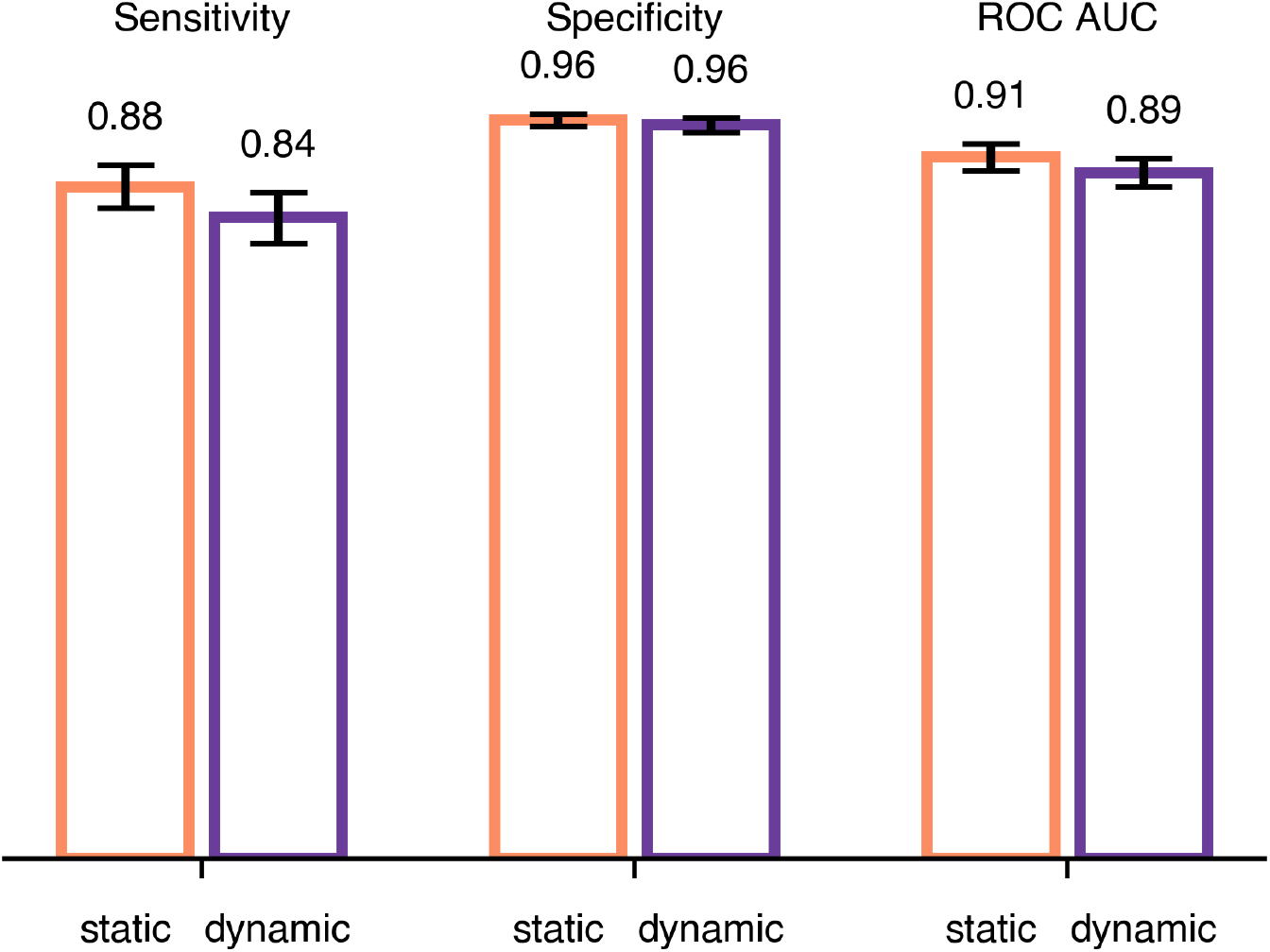
Mean sensitivity, specificity, and area under the ROC curve values for hyperparameter-tuned Decision Trees (DT) trained under a 50-repeat Monte Carlo cross-validation protocol with 80:20 training:validation splits. Standard error of the mean 95% confidence intervals are overlaid. The models were trained on SNAP2 score, and temperature factors, as well as either distance to rifampicin (‘static’) or minimum distance to rifampicin (‘dynamic’), and distance to mRNA (‘static’) or minimum distance to mRNA (‘dynamic’).

**Figure 6:**
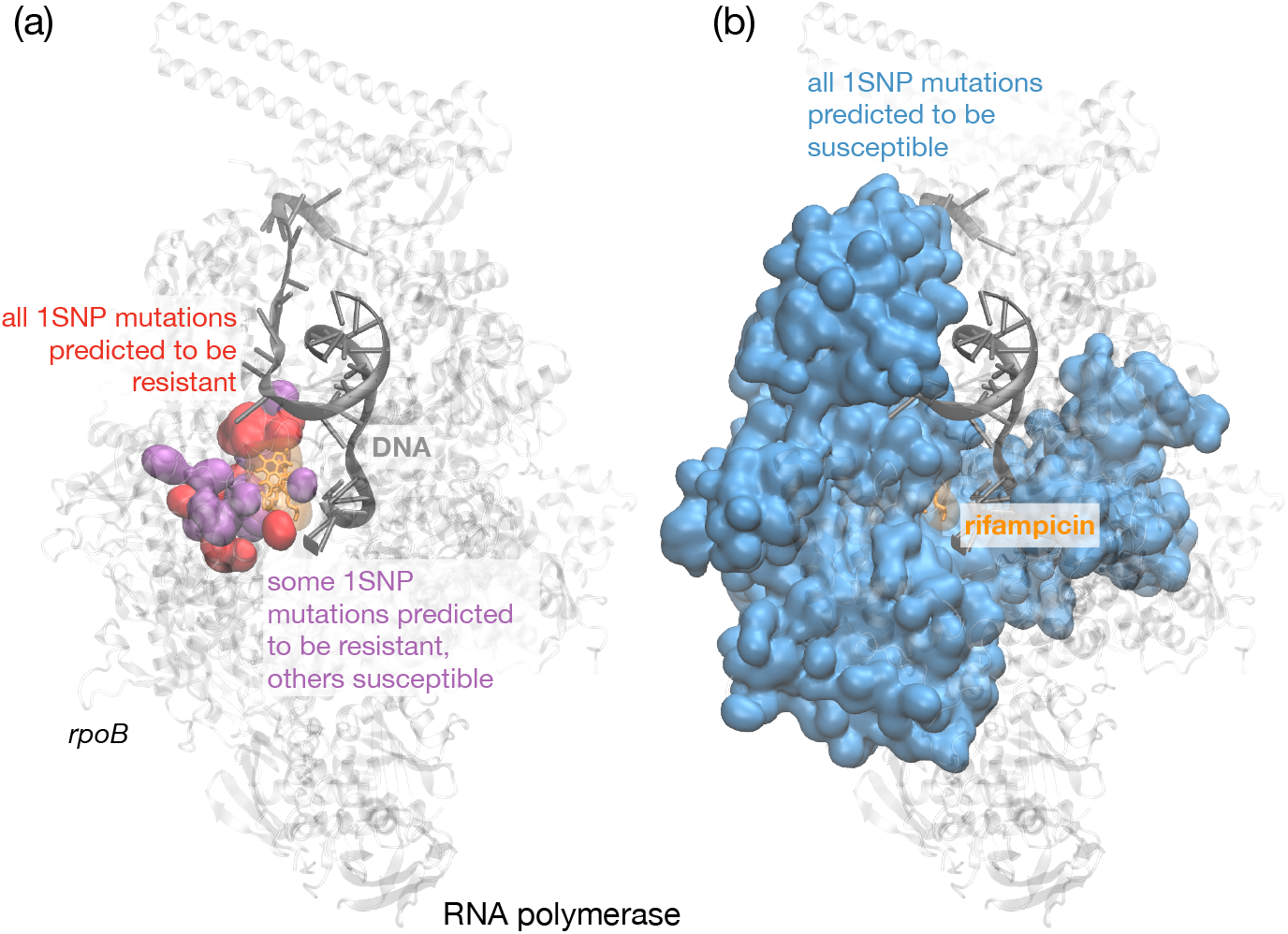
Mutations close to the rifampicin binding site are predicted to confer resistance. We used the ensemble of 50 decision tree models to predict all missense mutations in *rpoB*. (a) Close to the rifampicin binding site the models predicted that all mutations at a residue (red) led to resistance, whilst the mutation of some residues could lead to resistance or have no effect (purple). (b) Mutation of the majority of codons was predicted to have no effect, i.e. the bacterium would remain susceptible to rifampicin.

The ensemble of decision tree models consistently makes six very major errors (false negatives). These predominantly involve mutations located more than 13 Å from the rifampicin binding site that have been catalogued as resistant: these are E207K, L731P, S493R, V262A, S582A, and L378R. However, these mutations are very rare in the clinical dataset, with each mutation present in only two samples, suggesting that these errors may be attributable to laboratory mislabeling or phenotyping errors. The model also makes a number of major errors (false positives) within 13 Å of the rifampicin binding site, however the overall specificity is minimally affected due to the far higher number of correctly predicted susceptible mutations outside this region.

### Different models make different errors

If those six mutations were indeed a result of laboratory mislabelling, one might expect the other models to also misclassify them. However, by examining the mutations where at least one model consistently makes a mistake, we find that, despite all having similar performances, the ensembles of each of the four models all behave differently (Fig. S3). Closer inspection shows that the ensemble of logistic regression models makes several major errors (L443F, S493W/L, P454S), presumably because, unlike the tree-based methods, it cannot ameliorate the proximity of these residues to rifampicin.

### Dynamical features do not improve performance

Thus far all distances have been calculated from an experimental and therefore static structure of the RNA polymerase. *In vivo* and at physiological temperature the protein would be dynamic and it is notable that the temperature (*β*) factor, which includes some element of the protein dynamics, is one of the more predictive features. We therefore hypothesised that including some measure of the dynamics might improve the discrimination of the models. We measured the minimum distance between each residue and the centre of mass of rifampicin from a set of molecular dynamics simulations [37].

Training a decision tree, however, on the minimum distances to rifampicin and mRNA, the SNAP2 score, and temperature factors yielded no improvement over simply using the static distances, with a sensitivity of 0.835 *±* 0.033 and specificity of 0.956 *±* 0.009 (Table 1). With hindsight this is perhaps unsurprising, as although the magnitudes of the distances have changed, their magnitudes relative to one another remain fairly constant.

## Discussion

We can predict with moderate sensitivity whether individual mutations in the *rpoB* gene confer resistance to rifampicin using machine learning models trained on several structural, chemical and evolutionary features. As suggested by the spatial distribution of the mutations in the Train+Test dataset (Fig. 1b,c), any of the distances are moderately predictive on their own (Fig. 2a). SNAP2, which aims to predict the effect of mutations on a protein’s function, and the temperature factor are also predictive and were therefore included as features in our models. Because there are so few resistance-conferring mutations in the Train+Test dataset, and because one must ensure that any mutation in a Test dataset is not present in the Training dataset, it was not possible to use a conventional Train/Test split. Instead, we used Monte Carlo cross-validation with 50 repeats, with each iteration using a pre-defined random seed for reproducibility [26]. All the models performed similarly; we preferred the ensemble of decision tree (DT) models due to their simplicity and transparency. The DT models achieved, on average, a sensitivity of 0.876 ± 0.028 and a specificity of 0.963 ± 0.008. Interestingly, whilst all the models behaved similarly, they tended to misclassify different mutations. To investigate if including dynamic information improves performance, we replaced the distances observed from the static crystallographic structure [17] with the minimum distances measured from several classical molecular dynamics simulations but there was no significant effect. Including a measure of the protein dynamics will, we are sure, be important for some antibiotic targets.

To get an idea of how well our models performed compared to published models, we submitted our Train+Test dataset to the SUSPECT-RIF webserver [16]. The observed sensitivity of 0.935 was an improvement on the performance of our models which ranged 0.84-0.88 and a slight decrease on their reported value of 1.00. The specificity was, however, markedly different and we observed a value of 0.293 which is far lower than seen for our models (0.94-0.97) and also less than reported on their test set. This performance suggests, unfortunately, overfitting and also illustrates that, whilst recognising that reducing the very major error rate is the primary objective, one cannot neglect the major error rate if one is to train a model that could be suitable for clinical use. We also urge researchers to make their data and code publicly-available so models can be re-trained, results can be reproduced and model performance can be evaluated openly and fairly.

Whilst structure-based machine learning shows promise, it is most likely to succeed for drugs where the target gene is not essential as this typically leads to larger numbers of resistance- and susceptibility-associated mutations, such as pyrazinamide [14] and bedaquiline. We therefore expect similar problems to be encountered for the fluoroquinolones and isoniazid for tuberculosis where the target gene is essential. Alchemical free energy methods offer an alternative approach to machine learning; these use molecular simulation to calculate how the binding free energy of the antibiotic changes upon introduction of the mutation. These can be predictive [38, 39] but typically have large confidence limits [37] and require many orders of magnitude more computational resources than machine learning. These methods will therefore likely have to wait a few years until the amount of computational resources becomes feasible.

Our study has several shortcomings. By focusing on individual mutations we could only include samples with a single missense mutation in *rpoB* and some information will therefore have been discarded. This approach also precluded the possibility of including information from the other genes in the RNA polymerase: in theory this could boost predictive power by allowing the models to e.g. learn about compensatory mutations [40], but that is incompatible with the approach we used here. Choosing to use structural and chemical features of individual mutations also meant the models could only learn the effects of missense mutations: nonsense mutations, insertions, deletions and promoter mutations could not be predicted. One could, of course, allow other features, such as lineage, into the models. In theory this could improve performance, but in practice it is unlikely to have much of an effect due to the dominance of the distance-based metrics. We have also assumed that the protein structure is unaltered by the missense mutations. Since RNAP is essential, it is reasonable to assume any structural changes will be small, but it is likely that some of the mutations will lead to structural changes and this will not have been captured. As noted above, the paucity of resistance-associated variants introduces too much stochasticity into a Train/Test split and we caution that this could easily result in a “fortunate” separation of mutations leading to spuriously good performance; clearly both care and more data are needed. Finally, we have been conservative and reported performance at the level of individual mutations; this places undue weight on the rarer mutations and, if weighted according to the observed clinical prevalence, the calculated sensitivities would be higher and could be directly compared to those of the published catalogues [5, 6]. That said, this would not reflect the real world use case, which would be making predictions on samples containing mutations that are not present in resistance catalogues. This, of course, is challenging to assess since, by definition, those mutations are unlikely to be in any Train+Test dataset. Therefore, we chose to take the simplest approach and report all metrics at the level of individual mutations.

In addition to the usual plea for more data, especially samples with rifampicin minimum inhibitory concentrations which may improve model performance, we need to explore more sophisticated machine learning model approaches. In particular, we need to explore models that, rather than learning the effect of individual mutations, can learn the structural, chemical and evolutionary features of the full genetic sequence, ideally including all the genes that make up the RNA polymerase. This would maximise the information available to the models and would also allow nonsense mutations and insertions and deletions to be predicted (but still not promoter mutations). There is, however, no doubt that for genetics-based clinical microbiology to succeed, truly predictive methods, such as offered by machine learning, will be needed to fill the inferential gap when mutations not in any catalogue are encountered.

## Acknowledgemements

We are grateful for discussions with Professors Charlotte M. Deane, Timothy E. A. Peto and David Eyre.

## Reproducibility

The machine learning models can be trained and validated using the attendant online repository [41]. This also includes all source data and further code to regenerate (nearly) all figures in this paper.

