## Supplemental Information for "Predicting rifampicin resistance in *M. tuberculosis* using machine learning informed by protein structural and chemical features"

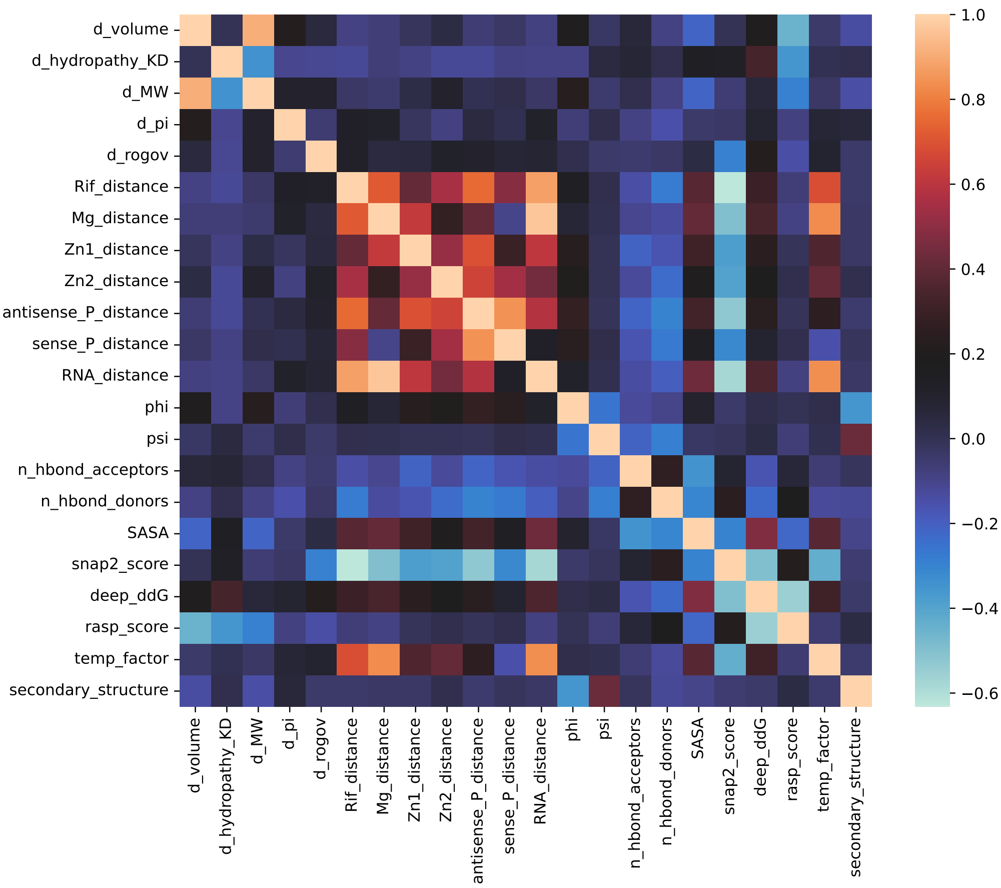


Figure S1: Pearson correlation coefficient matrix for each feature. Lighter colours represent feature value correlations.

*Current affiliation: Department of Biochemistry, University of Oxford, Oxford, UK
†These authors contributed equally.

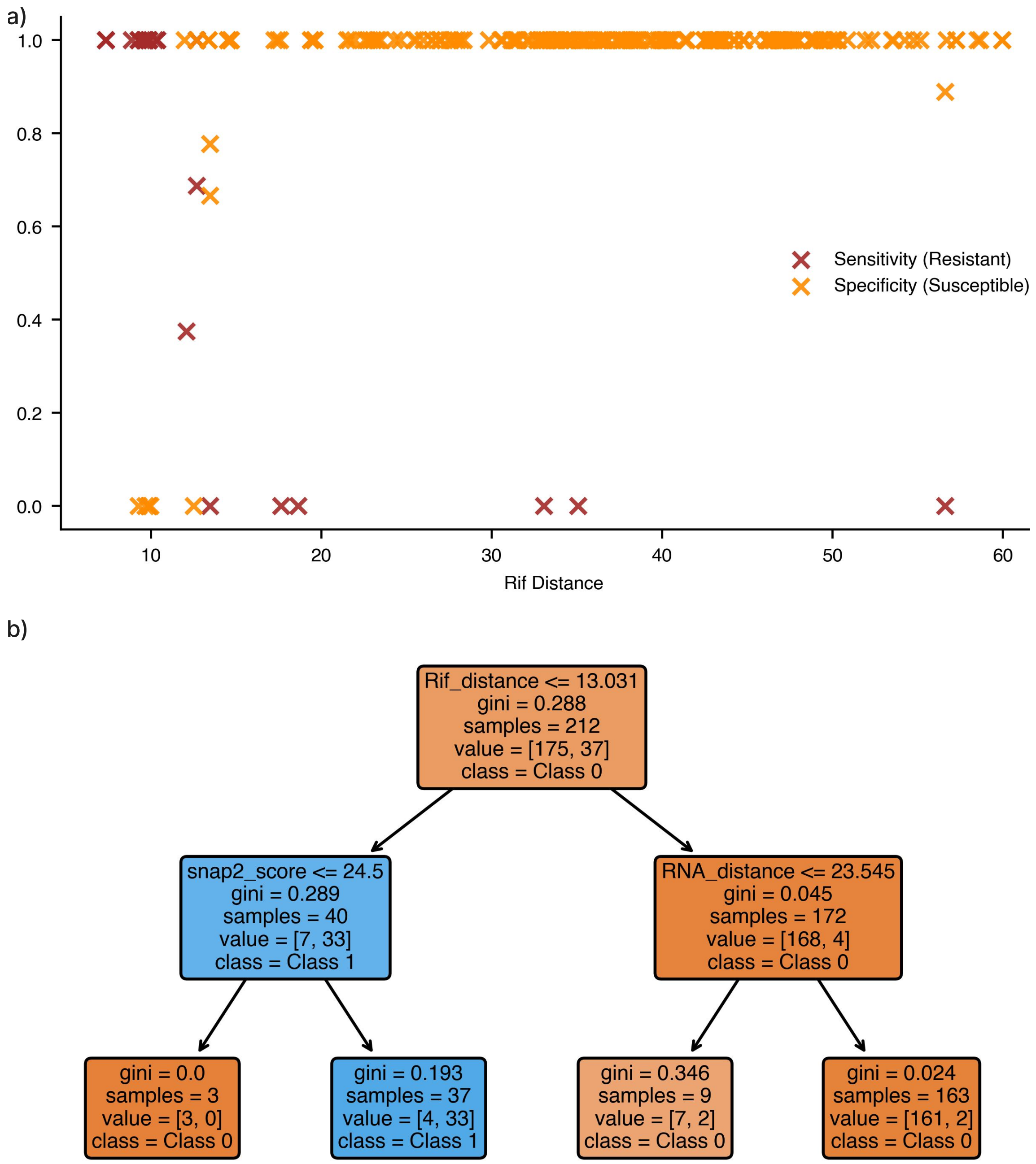


Figure S2: The Decision Tree models rely on distance from the rifampicin binding site. (a) How the sensitivity of resistant mutations (red) along with the specificity of susceptible mutations (orange) vary with the distance (Å) from the rifampicin (Rif) binding site as measured when part of the test set with a trained and hyperparameter tuned Decision Tree using 50 Monte Carlo cross-validation 80:20 splits. (b) An example of a Decision Tree trained on distance to rifampicin, distance to the mRNA, SNAP2 score, and temperature factor. The figure was generated by scikit-learn for one of the models trained during the cross-validation splits in Figure 4 of the main body of the paper.

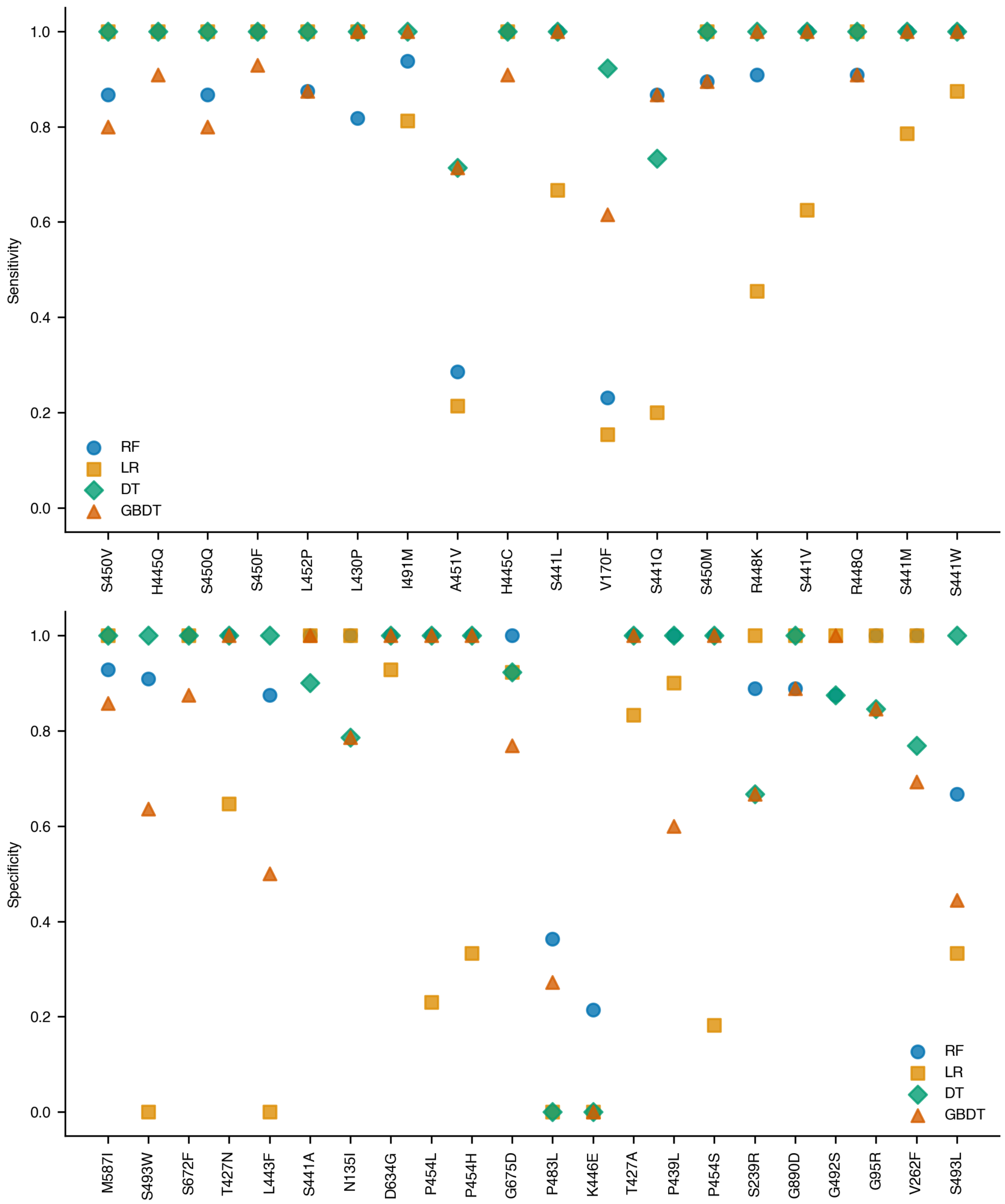


Figure S3: (a) Sensitivity and specificity of variants that are variably predicted by each model architecture, predicted under the same training and validation protocol as in Figure 4.
